# Intrinsically disordered proteins can behave as different polymers across their conformational ensemble

**DOI:** 10.1101/2024.09.27.615433

**Authors:** Saikat Chakraborty, Tatiana I. Morozova, Jean-Louis Barrat

## Abstract

Intrinsically disordered proteins (IDPs) are macromolecules, which in contrast to well-folded proteins, explore a large number of conformationally heterogeneous states. In this work, we investigate the conformational space of the disordered protein *β*-casein using Hamiltonian replica exchange atomistic molecular dynamics simulations in explicit water. The energy landscape contains a global minimum along with two shallow funnels. Employing static polymeric scaling laws separately for individual funnels, we find that they cannot be described by the same polymeric scaling exponent. Around the global minimum, the conformations are globular, whereas in the vicinity of local minima we recover coil-like scaling. To elucidate the implications of structural diversity on equilibrium dynamics, we initiate standard molecular dynamics simulations in the NVT ensemble with representative conformations from each funnel. Global and internal motions for different classes of trajectories show heterogeneous dynamics with globule to coil-like signatures. Thus, IDPs can behave as entirely different polymers in different regions of the conformational space.

**TOC Graphic:** 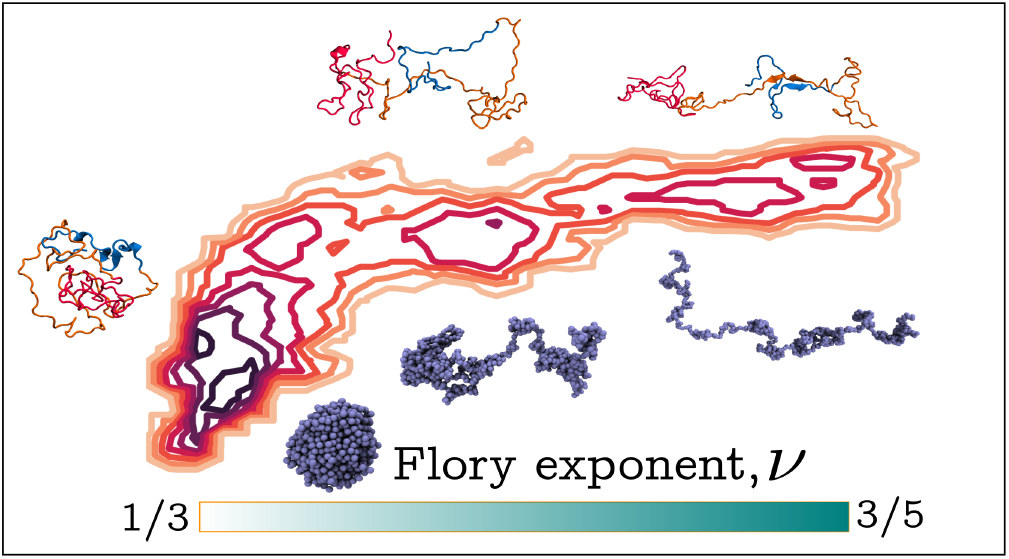

## Introduction

Proteins are complex biopolymers with amino acid (AA) residues being the building blocks or monomers. The conformations and equilibrium dynamics of the proteins are particularly sensitive to their AA sequences. Sequences resulting in native disordered three dimensional structures form the class of intrinsically disordered proteins (IDPs), which can explore a broad ensemble of configurations.^1^ The structural flexibility is key to their crucial role in control of genome size, binding to targets, and aggregations.^2^ The aberrant, fibrilar aggregations of some IDPs have also been implicated in several human diseases.^3^ Elucidating the relation between their structural diversity, and the equilibrium dynamics is crucial in understanding the functionality of this class of proteins.

The conformational heterogeneity of IDPs is reflected in a wide range and distribution of size, shape and compactness. In the realm of polymer theory,^4–6^ the IDP configurations are often characterized by the static scaling laws akin to globules, Gaussian random coil (RC), or self-avoiding walk (SAW).^7–10^ Single molecule spectroscopic measurements follow this approach through quantification of pairwise distances among the residues. ^8,11,12^ However, such a description becomes inadequate when multiple stable populations with drastically different structures exist in the free energy landscape.^13–15^ Particularly for polyampholytic IDPs there are theoretical predictions^13,16^ and numerical evidence^15^ of a two-state behavior and the bistability of structures. Even if some of the populations are rare for single IDPs, they may form a pathway to stable oligomers which lead to morphological diversity in aggregates.^17^

Assuming a specific class of polymer-like configurations implies well-defined scaling laws for the equilibrium dynamics, relating fluctuations of segments of different sizes,^4,5^ which can be used to analyze dynamic scattering experiments.^18,19^ When all the conformations are visited on a short time scale, of the order of the chain reorientation time, the situation is aptly represented by a statistical average corresponding to an equilibrium within this configurational ensemble. However, a different situation arises in the presence of multiple minima with energy barrier ∼ few *k*_*B*_*T*, that can prevent the barrier crossing for times comparable to the complete chain reorientation time. In this case, every protein can be observed in a separate minimum with distinct fluctuation dynamics, effectively behaving as a quasi stable conformer. Therefore, averaging over the conformational ensemble leads to a loss of information on the heterogeneity in structure and its implications for equilibrium dynamics.

Clearly, it is difficult to resolve the diversity in polymeric scaling of the quasi-stable conformers solely through experiments. Alternatively, atomistic molecular dynamics (MD) simulations grants a high resolution insight into individual populations.^15^ For instance, an observation of RC-like scaling in polyampholytic IDPs in single molecule experiments^14^ has been attributed to the bistability of globular and SAW-like structures using MD simulations.^15^ However, due to the presence of local minima, standard MD simulations may suffer from poor sampling of the energy landscape.^9,20–22^ Particularly, for long sequences the computational cost becomes prohibitive.

In this work, we perform explicit-solvent, all-atom molecular dynamics simulations with single chains of disordered protein bovine *β*-casein with 209 residues, having a molecular weight of ≈ 24 kDa.^23–25^ Apart from its importance in the dairy industry, this protein is speculated to display not only micelles, but fibrilar aggregation.^26–29^ Structural heterogeneity at the single molecule level often determines the aggregate morphology. ^17,30,31^ Therefore, we expect a rich conformational ensemble of this IDP. We sample the complete ensemble of conformations for *β*-casein using Hamiltonian replica exchange molecular dynamics (HREMD) simulations.^32–35^ To the best of our knowledge, this is the first time that the full ensemble is spanned for this naturally abundant protein.

Using the conformations obtained from this unbiased, enhanced sampling method, we obtain the corresponding free energy landscape (FEL). The FEL exhibits a global minimum, accompanied with two shallow and sparsely populated funnels. We probe the compactness of the conformations belonging to distinct funnels through scaling of internal distances among the residues with chain separation. Related Flory exponents show a wide variation, reflecting strong conformational heterogeneity. Structures around the global minimum are collapsed, whereas we observe more extended, coil like structures when conformers reside in local minima. Spectra of the segment specific fluctuation amplitudes also display consistent features for different conformational classes.

We elucidate the effect of structural heterogeneity on the equilibrium dynamics by initiating standard MD simulations in the NVT ensemble from representative conformations in each funnel. Probing the rotational and translational motion, we show that IDP can diffuse like a polymer globule or coil, depending on its location on the FEL. Internal fluctuations also display heterogeneity. For the trajectories around global minima, they are similar to those of a polymer globule, with whole body rotation being the primary mode of relaxation. In contrast, conformations in local minima have significantly larger internal fluctuations.

Micelle formation of *β*-casein is generally attributed to its block-copolymer like residue distribution.^24,25^ Three segments are usually defined with *N*-terminal spanning the first 40 residues, carrying a large net negative charge. The *C*-terminal with AA ∈ [136, 209] have many uncharged residues, resulting in strong hydrophobicity. The rest of the inner sequence (41 − 135) is characterized to be moderately hydrophobic. In this work, we also investigate the static and dynamic scaling features of each of these conventional segments. Through structural and dynamic characterization, we clearly distinguish the hydrophobic *C*-terminal as a collapsed structure with muted internal fluctuations. The other two segments can display swollen conformations similar to polymer coils.

## Methods

The 209 residue main chain sequence (PRO-0000004470) of entry P02666 (entry name CASB-BOVIN) on Uniprot database provides the AA sequence of *β*-casein. The structure predictor package I-TASSER ^36^ generates five initial configurations based on the sequence. We chose the conformation with the highest “confidence score” and performed all-atom MD simulations at the temperature *T* = 300 K in explicit solvent using GROMACS 2023 package. AMBER99SB-ILDN force field^37^ was utilized along with the TIP4P-D water model.^38,39^ The combination has been reported to be suitable to model IDPs.^40,41^ In our simulations, we consider neutral pH values which imply that the N- and C-termini of the IDP are charged (Arg1 – NH^+3^ and Val 209 – COO^−^, respectively).

We solvated the single protein chain in a cubic box with the initial size, *L*_*init*_ = 15.5 nm, which ensures that the chains do not interact with their periodic images. ^42^ First, the entire system is subjected to an energy minimization using the steepest descent method with the maximum force tolerance level set as 1000 kJmol^−1^nm^−1^. The energy of each system was minimized using 1000 steepest decent steps followed by a 1 ns equilibration at the NVT (number of particles, volume, and temperature) and the NPT (number of particles, pressure, and temperature) ensembles. Finally, the equilibration run in the NPT ensemble is performed for 200 ns.

All bonds were constrained using the LINCS algorithm. The Verlet leapfrog algorithm was used to numerically integrate the equation of motions with a time step of 2 fs. A cutoff of 1.2 nm was used for short-range electrostatic and Lennard-Jones interactions. Long-range electrostatic interactions were calculated by particle-mesh Ewald summation with a fourth-order interpolation and a grid spacing of 0.16 nm. The solute and the solvent were coupled separately to a temperature bath of 300 K using a velocity-rescaling thermostat with a relaxation time of 0.1 ps. The pressure was fixed at 1 bar using the Parrinello–Rahman algorithm with a relaxation time of 2 ps and isothermal compressibility of 4.5×10^−5^ bar^−1^.

We utilized the replica exchange with solute tempering 2 (REST2), employing the Hamiltonian replica exchange molecular dynamics (HREMD) simulation method, to augment the conformational sampling^33,34^ (See Figure S1 and S2 in the *Supplementary Information*). The REST2 method is integrated into GROMACS, patched with PLUMED.^43^ In this approach, interaction potentials within the protein (intraprotein) and those between the protein and the solvent are scaled by a factor λ and 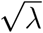, respectively. Whereas, water–water interactions remain unchanged. The scaling factor λ_*i*_ and the corresponding effective temperatures *T*_*i*_ for the *i*th replica are defined by the equations,

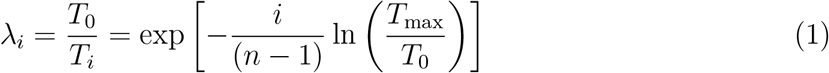

Here, *T*_0_ = 300 K and *T*_max_ = 450 K represent the effective temperatures of the lowest rank (unscaled) and the highest rank replicas, respectively, and *n* = 16 is the total number of replicas used. For analyses, only the trajectory of the unscaled lowest rank replica (λ_0_ = 1 or *T*_0_) is considered. Coordinate exchange between neighboring replicas is attempted every 400 MD steps. Simulations for each replica of HREMD spans 500 ns. The cumulative durations of HREMD simulations is 8 *μ*s.

For analyses of the equilibrium dynamics, selected conformations (see Results for the procedure of conformation selection) were subjected to MD simulations in NVT ensemble. This avoids artefacts in calculations of mean squared displacements due to fluctuations in the size of the periodic cubic box. ^44^ The protocol in the simulation remains same as the equilibration NPT simulation. However, the pressure coupling is turned off during this simulation.

We analyze the conformations and the trajectories with home grown Python scripts and in-built programs of GROMACS. ^45^ The secondary structure prediction was computed using DSSP. Alongside, the MDAnalysis software have been used for reading trajectory files and analyzing the data.^46^ We quote the statistical errors on the fit parameters from the Jackknife analyses with each Jackknife bin containing all but one sample. ^47^

## Results

### Conformational Heterogeneity of the IDP

#### Free energy landscape of *β*-casein

A commonly used reaction coordinate to identify heterogeneity in sizes of the IDPs is the radius of gyration,

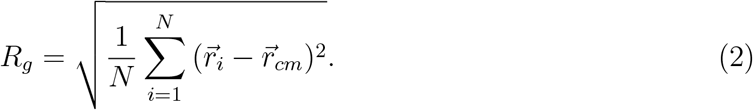

Here, 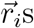 are the coordinates of *N* = 209 C_*α*_ atoms of *β*-casein. 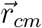 denotes the positions of the centre of mass,

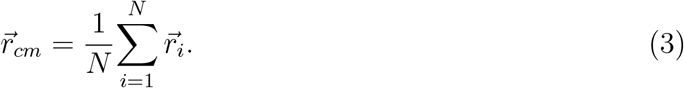

Also, the estimate of the solvent-accessible surface area (SASA), *S*_*s*_ reflects the exposed area of the protein to the solvent. Thus, we begin with the presentation of the free energy landscape (FEL) of the IDP as a function of *R*_*g*_ and SASA, see Fig. 1. We draw the FEL computing the joint probability *P*(*R*_*g,α*_, *S*_*s,α*_) from histograms of the variables *R*_*g*_ and *S*_*s*_.^48^ We use HREMD simulations^32–35^ to ensure unbiased enhanced sampling of the conformations. Our simulations employ AMBER99SB−ILDN force field along with the TIP4P-D water model.^37,38^ The combination has been reported to be suitable for the simulations of the IDPs.^40,41^ Consistency between the small angle X-ray scattering (SAXS) profiles obtained from the experiments^25^ and that from the HREMD generated conformations (see Figure S3 in *Supplementary Information*) over a wide range of *q* values indicates the efficiency of the sampling.

**Figure 1:**
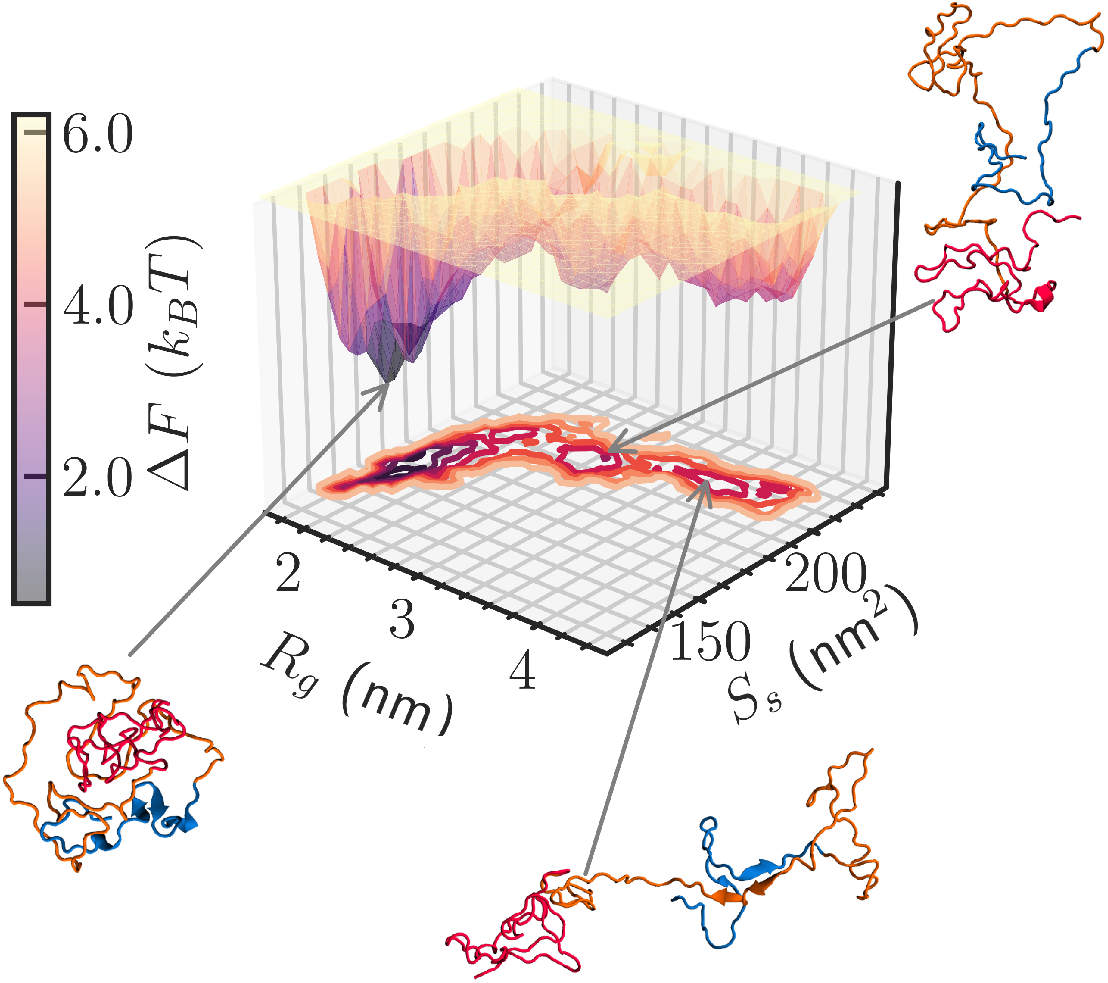
Free energy landscape of *β*-casein. Free energy landscape of *β*-casein in the radius of gyration (*R*_*g*_)-SASA (*S*_*s*_) plane, and its three-dimensional representation with the depth indicating the differences in free energy. A global minimum with two shallow funnels are observed. The arrows indicate the minima to which the representative configurations belong. Distinct colors mark commonly differentiated segments of block copolymer-like *β*- casein: Hydrophilic *C*-terminal (AA 1−40) is colored blue. Orange and red regions discern inner moderately hydrophobic (AA 41−135) and hydrophobic *N*-terminal (AA 136−209). The largest conformations have a strong tendency to develop *β*-sheet between the first two segments.

The distance between two contour lines on the *R*_*g*_ − *S*_*s*_ plane provides an estimate of the free energy difference between two states *α* and *β* following the equation,^48^

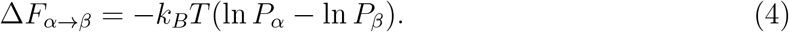

The three-dimensional energy landscape displays a deep funnel, indicating a global minimum, and two relatively shallow funnels. The latter two contains about 10% and 8% of the population, respectively. Moreover, multiple structures separated by small energy barriers are present in every funnel. This is a characteristic feature of the IDPs. ^49–51^

The configurations populating the lowest free energy minimum are compact, as may be noticed already by a visual assessment of the conformations. Local minima, on the other hand, contain extended structures. Upon closer inspection, we find that the hydrophobic *C*-terminal (colored red) is nearly invariably collapsed. Whereas, the inner, moderately hydrophobic domain (colored orange), and the hydrophilic *N*-terminal (colored blue) have varying compactness for various funnels. Furthermore, the conformations with large *R*_*g*_ values typically form *β*-sheets between the protein’s first two segments.

#### Characterization of the conformations: Global and local observables

The probability density distribution of *R*_*g*_, *P*(*R*_*g*_) shows that *β*-casein conformations densely populate the region with *R*_*g*_ ≤ 2.5 nm, cf. Fig. 2a. The region corresponds to the funnel with the global minimum in the FEL. We assign a label C to these compact conformations. On the other hand, *R*_*g*_ > 3.4 nm characterizes the most extended and sparse population (label E). For the intermediate values of *R*_*g*_ ∈ [2.5, 3.4] nm, another sparsely populated region can be identified. These conformations have intermediate sizes and we will refer to them as class I.

**Figure 2:**
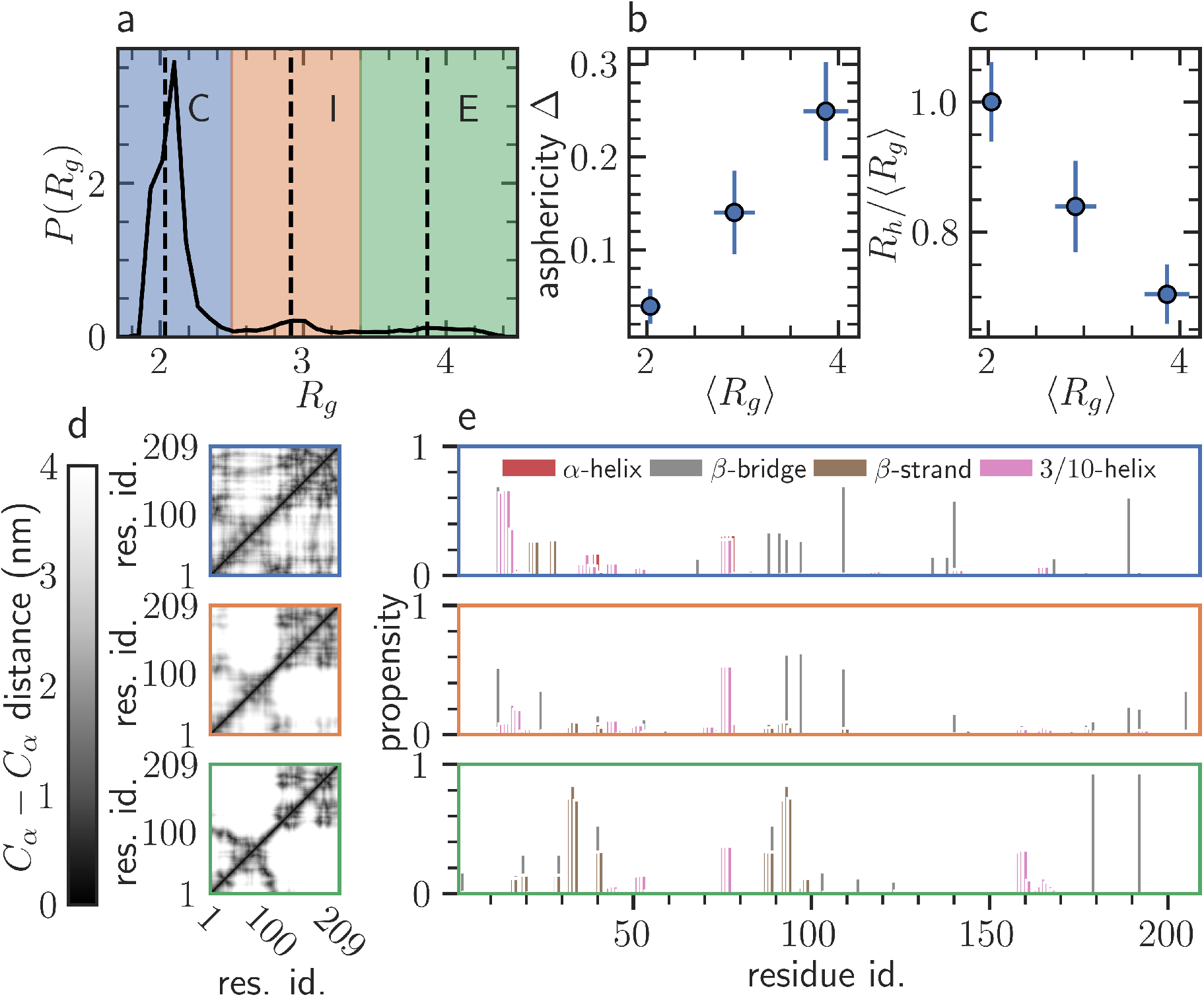
Discerning different classes of conformations. (a) Probability density distri- bution of *R*_*g*_, *P*(*R*_*g*_) shows densely populated blue marked region with label C. Two other ranges of *R*_*g*_, colored orange (label I) and green (label E), correspond to sparse population of conformations. Dashed vertical lines indicate the average values of *R*_*g*_ (⟨*R*_*g*_⟩) for each class of conformations. Plots of (b) asphericity, Δ, and (c) hydrodynamic ratio, *R*_*h*_/⟨*R*_*g*_⟩ versus ⟨*R*_*g*_⟩. Microscopic differences in different classes as revealed by the (d) average pair- wise distances between residues (top to bottom: C, I, and E), and (e) secondary structure propensities (top to bottom: C, I, and E).

We measure the deviation from spherical symmetry of the conformations using the asphericity,^52–54^

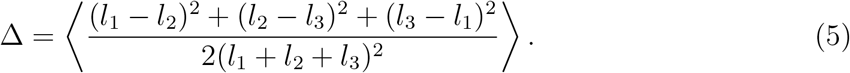

Here, *l*_1_, *l*_2_, and *l*_3_ (*l*_1_ > *l*_2_ > *l*_3_) are the eigenvalues of the gyration tensor. Here and below, the notation ⟨..⟩ denotes averaging over the conformations within a specific configurational class. Fig. 2b shows Δ for ⟨*R*_*g*_⟩ corresponding to the three different classes. C-type conformations with Δ ≈ 0 are almost spherically symmetric. Δ increases monotonically with ⟨*R*_*g*_⟩ with value for the class E being close to that of a three dimensional random walk with Δ ≈ 0.38.^55^

Furthermore, the three classes differ significantly in their values of the hydrodynamic ratio, *i*.*e*., the ratio between hydrodynamic radius, *R*_*h*_ and *R*_*g*_ (Fig. 2c). For a particular conformation, we determine *R*_*h*_ using the Kirkwood approximation that treats the intramonomer hydrodynamic interactions with the Oseen tensor. ^56^ The approximation reduces the calculation of *R*_*h*_ to the reciprocal of the average of the inverse of the distances between all pairs of C_*α*_ atoms *i* and *j, d*_*ij*_, *i*.*e*.,^41,56–59^

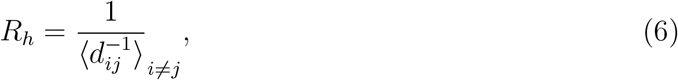

E type conformations show *R*_*h*_/⟨*R*_*g*_⟩ ≈ 0.7, similar to that for a Gaussian chain (*R*_*h*_/*R*_*g*_ ≈ 0.7).^5^ On the other hand, for structures belonging to class C, the ratio is closer to the hard sphere limit 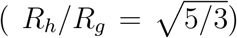.^5^ The I type conformations exhibit intermediate values of *R*_*h*_/⟨*R*_*g*_⟩.

Differences among the different types of conformations are evident in the average pairwise distances between residues as well, see Fig. 2d. Residues in different segments of the chain are close to each other in C types of conformations, indicating a compact structure. However, the hydrophilic *N*-terminal and hydrophobic *C*-terminal are well separated for class E. Once again, the structures belonging to class I show intermediary features in regards to the proximity between residue pairs that are neither as homogeneous as C type nor as separated as E type conformations. For all conformational classes, majority of secondary structures develop in the *N*-terminal and the inner domain (Fig. 2e). Thus, the compactness of the *C*-terminal across the structural ensemble is predominantly driven by hydrophobic collapse. Also, most of the helical structures for the majority C type population appear within the first 40 residues of the protein. ^25^ Moreover, between these first two segments of *β*-casein, the E type conformations show strong propensity to form *β*-sheets.

The E and I type conformations are relatively infrequent (18%) in the ensemble of the IDP conformations. However, such populations are transient states that have been linked to diversity in aggregate morphology. ^60,61^ Particularly, the *β*-sheet rich E conformations can result in the speculated fibrilar aggregates of *β*-casein.^27^ Also, the conformations around different funnels exhibit spherical to extended structural signatures, which may give rise to different polymeric scaling behaviors. Therefore, we analyzethese static and dynamic scaling properties separately for different conformations.

### Static polymer scaling

#### Compactness of the whole chain and its blocks

For complex biopolymers such as proteins, the average internal distances, *d*_*ij*_, can capture the compactness of the conformations. Polymer theory predicts a power-law profile, ^4,5^

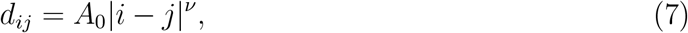

where *A*_0_ is a constant prefactor and *ν* is the Flory exponent. When intraprotein attractions overpower the solvent-protein interactions, a collapsed structure is expected with *ν* = 1/3. If the solvent-protein interaction dominates, proteins have *ν* ≈ 0.6 as a SAW. For comparable relative strengths of these two interactions, the protein behaves like a Gaussian random coil, being characterized by *ν* = 1/2. We evaluate these scaling features plotting ⟨*d*_*ij*_⟩ versus chain separation |*i* − *j*| on double-log scales for different types of conformations, see Fig. 3a. For small |*i* − *j*|, *ν* ≈ 1 because of the stiffness of the protein. Measurements of *d*_*ij*_s for large |*i* − *j*| values suffer from poor sampling. Thus, for obtaining *ν* values we fit the data points at intermediate values of |*i* − *j*| to Eq. (7). The estimates are quoted in Table 1.

**Table 1:**
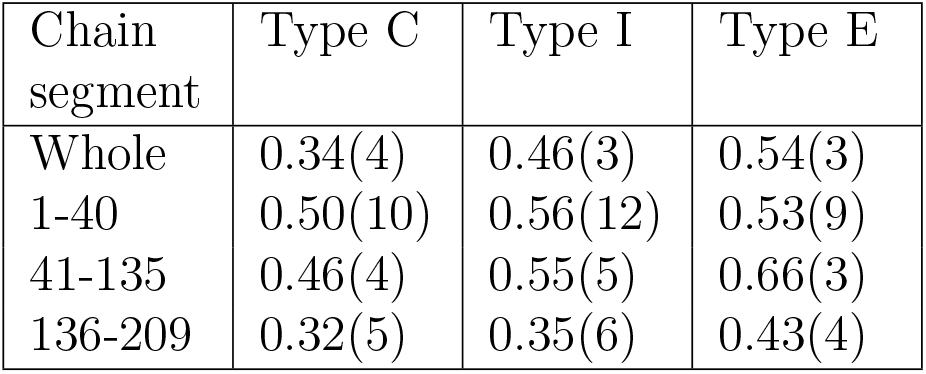
Estimates of Flory exponents. Calculated values of *ν* for the entire chain and conventional segments of *β*-casein. Results are obtained by fitting the scaling relation in Eq. (7) with the data in Fig. 3. Statistical errors are estimated using the Jackknife error analysis.

| Chain segment | Type C | Type I | Type E |
| --- | --- | --- | --- |
| Whole | 0.34(4) | 0.46(3) | 0.54(3) |
| 1-40 | 0.50(10) | 0.56(12) | 0.53(9) |
| 41-135 | 0.46(4) | 0.55(5) | 0.66(3) |
| 136-209 | 0.32(5) | 0.35(6) | 0.43(4) |

**Figure 3:**
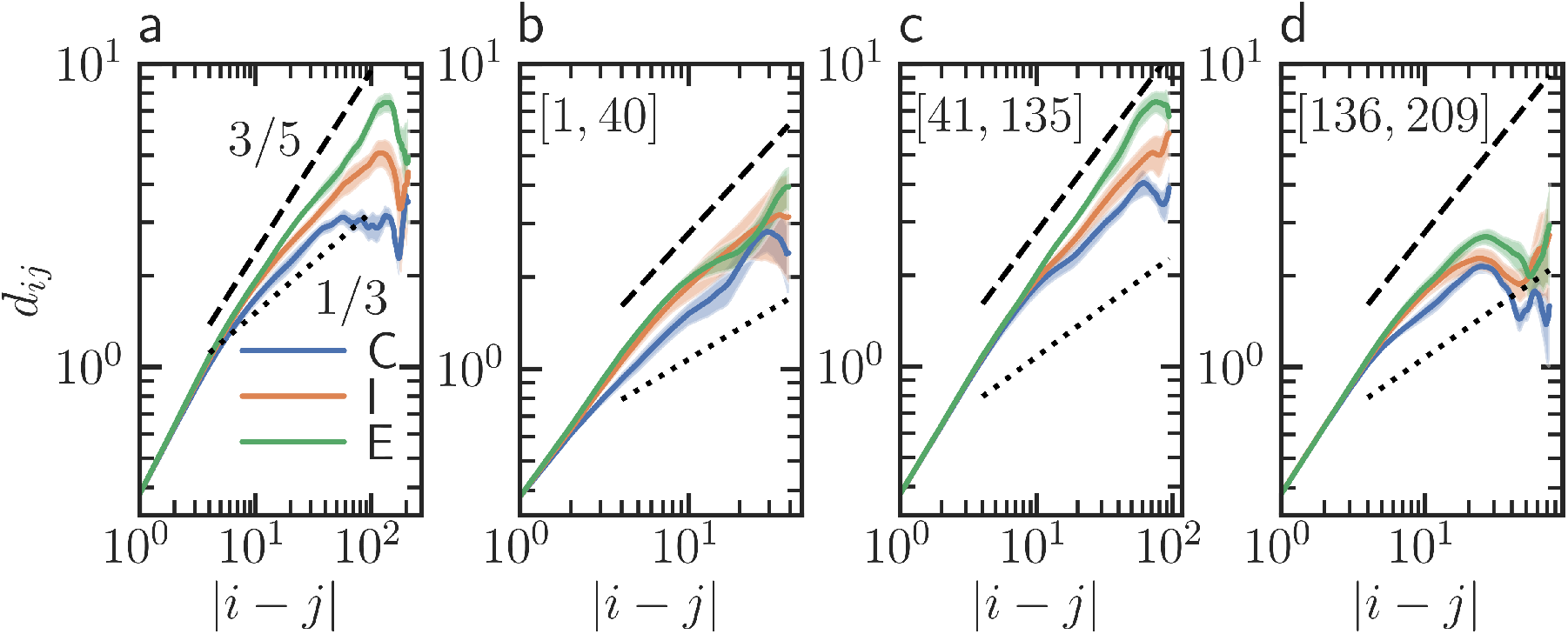
Compactness of the chain and its segments. (a) Log-log plot of internal distances *d*_*ij*_ as a function of the chain separation |*i* − *j*| between residues with indices *i* and *j*. Results are for the entire chain and three different classes of conformations. Dotted and dashed lines are power laws with quoted exponents. They refer to scaling of collapsed structure and SAW. (b-d) Same as (a), but for different segments mentioned on each frame.

Conformations belonging to group C exhibit scaling similar to the one of a collapsed polymer. In contrast, those with label E on average have *ν* ≈ 0.54 indicating extended structure. Consistent with observations above, structures belonging to class I display an intermediate value of *ν*. The wide variety in the values of *ν* for different populations indicates that IDPs can behave as a different polymer in different parts of the ensemble.

The values of *ν* for the whole chain describe the degree of compactness of the conformations in different funnels. However, the estimates may be effective exponents, originating from combinations of drastically different scaling behaviors of its three chemically distinct segments. Thus, we proceed to investigate the Flory exponents of the three segments of *β*-casein, see Fig. 3(b-d). In all conformational types, the hydrophobic *C*-terminal is observed to be close to collapsed states. The *N*-terminal, consisting of 40 residues, exhibits scaling that is approximately Gaussian coil-like across all conformations. Significant differences in the scaling behavior are evident in the inner moderately hydrophobic domain, which spans 94 residues. For C type conformations, the *ν* value falls between those of the N and *C*-terminals, indicating a slightly collapsed Gaussian coil. In class I configurations, scaling behavior of this domain and that of hydrophilic *N*-terminal are similar. Conversely, in class E configurations, the inner segment appears more extended than a polymer coil with volume exclusion effects. This phenomenon may arise from the restricted dihedral space of an AA sequence, which enhances the effective excluded volume. This effect and the stabilization of extended conformation of the domain due to formation of the *β*-sheets with the *C*-terminal may result in this high value of *ν*. Notably, it is difficult to obtain such detailed insights into the internal structures of chemically distinct segments through dynamic scattering, or single molecular experiments.

#### Scaling of Rouse mode amplitudes

Conventionally the Rouse model is used to describe the dynamics of IDPs.^62,63^ However, the spectral profiles of the Rouse mode amplitudes can be an alternate probe to analyze the internal structures of the IDP conformations.^64^ The Rouse model represents a polymer with *N* beads connected by harmonic springs having mean squared bond length *b*^2^. The effect of solvent is implicitly accounted for introducing a monomer friction coefficient and a random force. The resulting 3*N* coupled differential equations can be decoupled *via* orthogonal transformation of the monomer coordinates **r**_*n*_ to Rouse modes *X*_*p*_(*t*), *p* = 0, 1, .., *N*. Modes with *p* > 0 relates to average internal fluctuations of a segments of *N*/*p* monomers. A closed form can be obtained for the mean-squared mode amplitude, 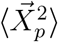 of the *p*th mode that show scaling of the form,^4,5^

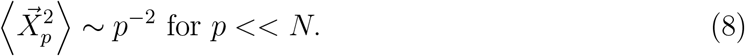

This theoretical framework was developed for ideal RW-like polymers. Nevertheless, physical volume of monomers in real chains can be introduced, mapping it to a SAW using the Flory exponent. The scaling in Eq. (8) thus modifies to^64,65^

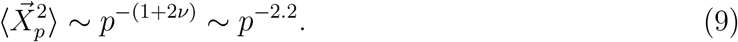

Notably, for higher values of *ν* the power-law decay can be faster. For the large *p* or small length scales excluded volume dictates shape fluctuations and scaling of the form of Eq. (9) is expected to hold. Whereas, effects of intrachain interactions becomes important in longer segments which recast Eq. (9) to 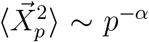 for small *p*. In the context of coil to globule transition of polymers, it has been recently shown that the mode spectra displays continuous variation in *α* for different *ν* values, decided by the effective strength of interactions.^64^ The spectral profile for the limiting case of collapsed structure or *ν* = 1/3 can be obtained by describing globule structures as ideal polymer in a spherical confinement of radius *R*_*g*_.^66^ Monomers in two sufficiently long subchains (therefore small *p*) are uncorrelated. Therefore, collapsed conformations correspond to *α* ∼ 0. The upper bound of *α* = 2.2 is set by the SAW.

The wide variation in *ν* values across different conformation classes (Fig. 3, Table 1, and related text) implies variation in power-law decay profile of 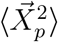. We analyze the diverse profiles for the entire chain (Fig. 4a) as well as for three specific segments (Fig. 4(b-d)). To facilitate a comparison of segments with different number of residues, *N*_*seg*_, we present results as a function of *p*/*N*_*seg*_ (*N*_*seg*_ = *N* for the whole chain). For all the frames, data for *p*/*N*_*seg*_ > 1/3 represent subchains containing fewer than three C_*α*_ atoms, incorporating the effects of bond angles and fluctuations. Notably, within the range *p*/*N*_*seg*_ ∈ [1/5, 1/3] show faster than 2.2 decay. This is due to the chain stiffness. As observed in Fig. 3, differences in effective elasticity of the whole chain and its different segments result in variation in the lower range values.

**Figure 4:**
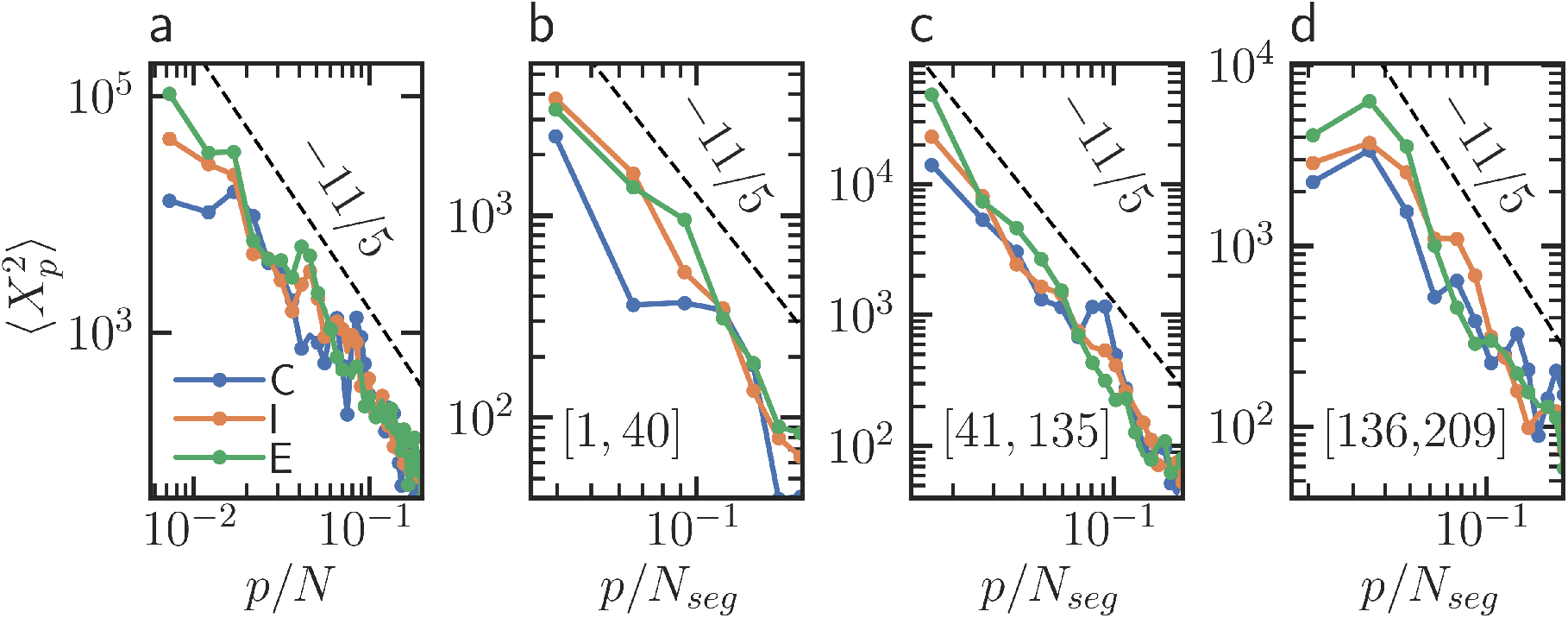
Spectra of the Rouse mode amplitudes. (a) Profile of mode amplitudes for the three classes of conformations. The dashed line indicates a power-law decay with an exponent − 2.2. The results are for the whole chain. (b-d) Same as (a), but for the segments with residue window mentioned inside the frames. We have smoothed the data through a moving average over neighboring pair of data points.

The spectra at lower *p* values characterize the conformations; however, this region is linked to long subchains that exhibit poor sampling. Consequently, we can only provide qualitative insights into the structural differences. In Fig. 4a, the C type conformations display relatively flat spectra for the initial modes, indicative of a globular structure. Distinguishing between the I and E class conformations in this region proves challenging.

For the hydrophilic *N*-terminal (Fig. 4b) and inner segment (Fig. 4c), proximity to *p*^−2.2^ profile indicates relatively extended structures. Conversely, in all classes of conformations, spectra for hydrophobic *C*-terminal (Fig. 4d) are nearly independent of *p*. This suggests collapse of the segments across various conformation types. The trends in scaling features are in line with the structural information obtained from the scaling of the form of Eq. (3).

### Effect of structural heterogeneity on fluctuation dynamics

#### Translational and rotational diffusion

In order to investigate the effects of multiple energy minima on equilibrium dynamics, we analyze the temporal evolution of configurations associated with these funnels. We initiate molecular dynamics (MD) simulations in the NVT ensemble, starting from random configurations across three distinct classes, and monitor their radius of gyration (*R*_*g*_).

The conformations of *β*-casein exhibit intricate intra-protein interactions, such as changes in secondary structure propensities and the formation and annihilation of hydrogen bonds. These fluctuations result in conformational switches among the three configurational classes. We use the preassigned ranges of *R*_*g*_ corresponding to the three types of conformations to select sections of trajectory that stay within a specific class. Thus, we retain the labels C, I, and E for the trajectories of conformational fluctuations. We present results from 5 such trajectories, each being 80 ns long, from each classes of conformations. The quoted time span is comparable to that of the slowest mode of relaxation for the conformations, see Figure S4 in *Supplementary Information*.

The translational diffusion of a solute at temperature *T* in a solvent with viscosity η is conventionally described by the Stokes-Einstein-Sutherland equation,^4,5^

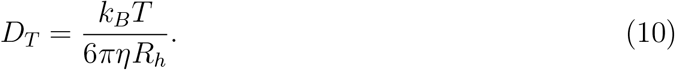

Here *D*_*T*_ is the translational diffusion coefficient, and *k*_*B*_ is the Boltzmann constant.

To extract *D*_*T*_ we estimate the time averaged mean squared displacements (MSDs), 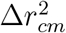 of the COM of the protein at time *t*,

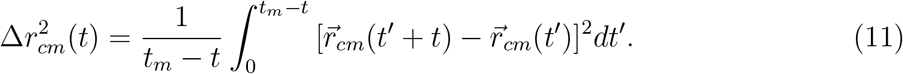

Here *t*_*m*_ = 80 ns is the time of measurement.

In Fig. 5a, we plot the time dependence of 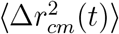 for different classes of conformations. It is difficult to separate different classes. This is likely due to lack of sampling and the relatively short time window that hinders appropriate time averaging. Moreover, *D*_*T*_ is inversely proportional to *R*_*h*_ (Eq. (10)). Earlier static estimates (Fig. 2c) show that 1/*R*_*h*_ only decreases by ≈ 25% when conformations moves from class C to E. We estimate diffusion coefficients *D*_*T,PBC*_ fitting the MSDs to the expression,

**Figure 5:**
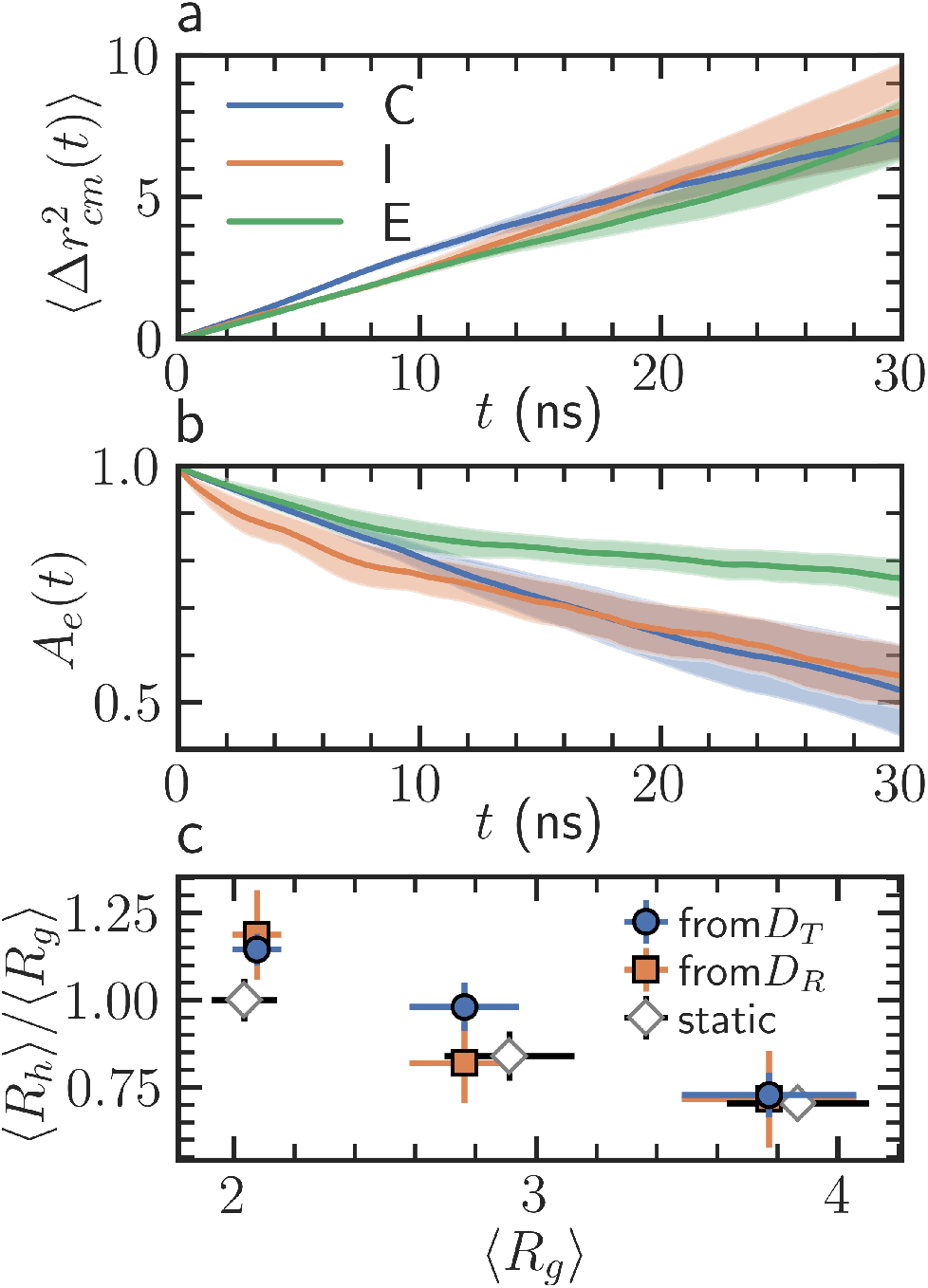
Translational and rotational diffusion. (a) MSD of the center of mass, 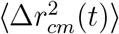 for three different classes of conformations. Shaded region indicates standard error. (b) End-to-end vector autocorrelation, *A*_*e*_(*t*) corresponding to trajectories in different funnels. (c) Hydrodynamic ratio, *R*_*h*_/*R*_*g*_ for fluctuations of C, I, and E types conformations. The estimates are obtained from the fitting of the early time data in (a), and (b). Compar- ison of the dynamic estimates with the static ones obtained from 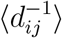.

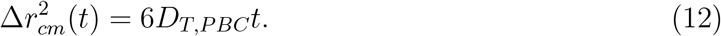

For long *t* statistical fluctuations are stronger in the data. Therefore, the fitting is performed for *t* ≤ 20 ns. Such procedures on trajectories generated with MD simulations with periodic boundary conditions (PBC) underestimate the value of the diffusion coefficient which is corrected following,^57,67^

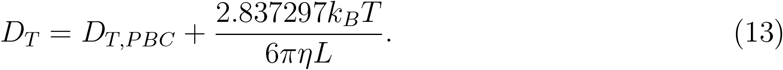

The measurement of *D*_*T*_ leads to evaluation of *R*_*h*_ through Eq. (10). Fig. 5c illustrates the corresponding *R*_*h*_/⟨*R*_*g*_⟩ in relation to the ratio ⟨*R*_*g*_⟩ for different conformational classes.

The rotational diffusion coefficient, *D*_*R*_, of a solute is given as,

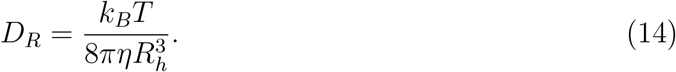

In our simulations with PBC, we compute the rotational diffusion coefficients, *D*_*R,PBC*_ from the time averaged autocorrelation, *A*_*e*_(*t*), of the end-to-end unit vector between two C_*α*_ atoms of the residues at the chain ends, 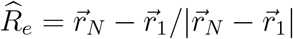,

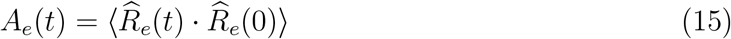

Plots of *A*_*e*_(*t*) display different decay rates for the three classes of trajectories, see Fig. 5b. Estimates of *D*_*R,PBC*_ are obtained from these correlation functions using the expression,

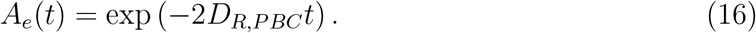

Though less pronounced than its translational counterpart, the under evaluations of rotational diffusion coefficients due to the finite size effects are corrected following,^68^

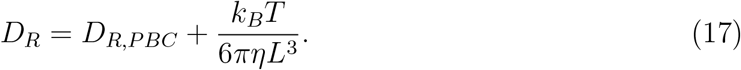

From the estimates of *D*_*R*_ we extract *R*_*h*_ using Eq. (14) and demonstrate the variation of *R*_*h*_/⟨*R*_*g*_⟩ for different classes of trajectories, see Fig. 5c. The values of the ratio closely align with those obtained from *D*_*T*_, and are also globally consistent with those obtained from the static calculation using the Kirkwood’s approximation. Note that for SAW-like polymers, it is known that the Kirkwood approximation (Eq. (6)) overestimates *R*_*h*_ by less than 4%.^58^ The different values of hydrodynamic ratio suggests that, in each of the different classes and at short time scales, the protein behaves akin to different kinds of polymers, going from globular to SAW behaviour as *R*_*g*_ increases (see also the discussion following equation (6)). Experiments, in which this ratio can be determined using a static approach (e.g. single molecule spectroscopy experiments^8^) or a dynamical one (e.g. inelastic scattering experiments) would in general lead to a weighted average of these different behaviors.

In spite of the good agreement between static and dynamic determinations, we note that for C conformational classes (smallest ⟨*R*_*g*_⟩), the values of *R*_*h*_/*R*_*g*_ obtained from analyses of equilibrium dynamics appear to be slightly larger than the static one. The mismatch may originate from extraction of *D*_*T*_ or *D*_*R*_ from a short time window. For instance, C type conformation appear to be more compact in dynamic measurements. A time window in which relatively flexible segments of the protein do not relax completely can result in such an apparently higher value of *R*_*h*_/*R*_*g*_. Poor sampling of the conformational space may also cause the deviation.

#### Relaxation of the Rouse modes

While insights on global dynamics of the protein are obtained from *D*_*T*_, and *D*_*R*_, internal fluctuations at different scales can be elucidated probing the relaxation of the Rouse modes, 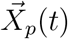. For an ideal RW-like polymer, the Rouse model predicts an exponential decay for the autocorrelation of 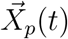. The characteristic relaxation time, *τ*_*p*_ follows a power-law decay of the relaxation time,^4,5^

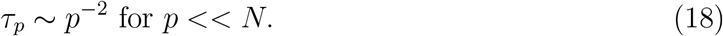

In addition, monomers in dilute solution drags the surrounding solvent with it. This results in a flow that applies force on other monomers in the polymer. Such hydrodynamic interactions are not considered in the Rouse model. The interaction reduces the separation among the relaxation time for different modes. The resulting spectrum is described by the Zimm model as,^4,5,69^

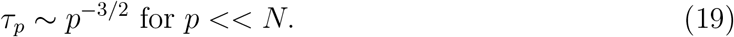

The spectra of *τ*_*p*_ for the whole chain vary widely for different conformational classes, see Fig. 6a. For convenient comparison of the spectra for different trajectory classes, the vertical axis is normalized by *τ*_1_, which corresponds to the slowest mode of relaxation of the protein. Also, scaling the *p* by *N*_*seg*_ facilitates relative analyses of segments with different number of residues. The C type conformations show almost no mode separation in timescales. Such a flat spectra indicates a compact, solid like structure. Moreover, The *τ*_*p*_s in this case are comparable to the decorrelation time of *A*_*e*_(*t*), see Figure S4 in *Supplementary Information*. Therefore, global rotation is the primary mode of relaxation for C type conformations. A slightly steeper spectra, suggestive of higher degree of internal fluctuations, is observed for equilibrium fluctuations of the class I. Such profile is consistent with previously noted reduced compactness in conformations, which exhibit diminished contact among the residues. Most prominent *p* dependence of *τ*_*p*_ can be noted for fluctuations of E type conformations. Therefore, significant degree of heterogeneity in internal fluctuation dynamics exits for different conformations of the same IDP.

**Figure 6:**
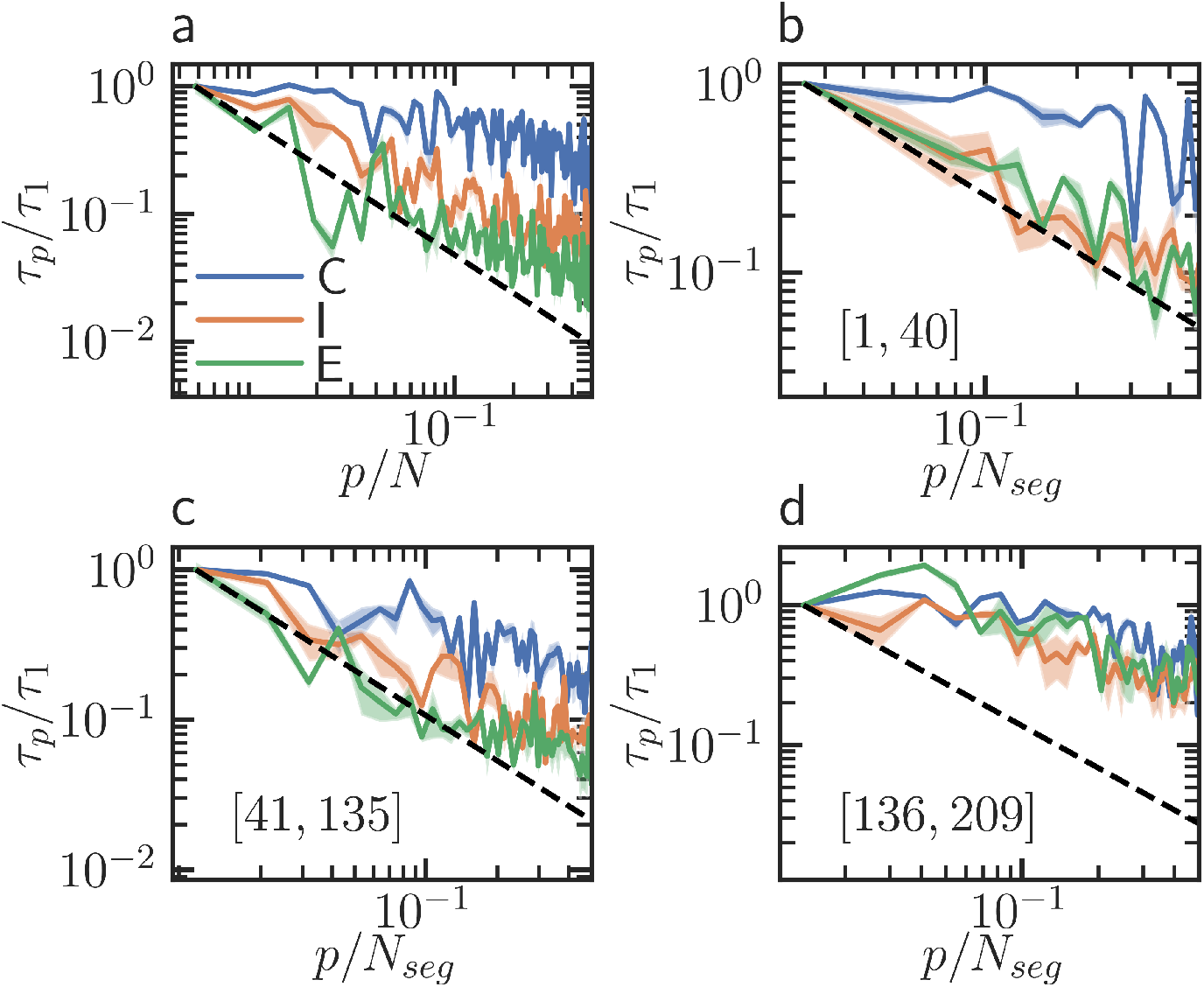
Spectra of relaxation time. Relaxation time, *τ*_*p*_, associated with the *p*th mode, scaled by *τ*_1_. For the whole chain (a), and the three segments mentioned in the frames (b-d). The dashed line indicates a (*p*/*N*_*seg*_)^−1^ decay for reference. Note that, this line indicates a significantly slower decay than Zimm model predictions (see text for details).

Though conformational class E displays a static scaling similar to a swollen coil, the profile of *τ*_*p*_ exhibit slower than *p*^−3*/*2^ decay, predicted by the Zimm model.^69^ In polymeric representation of protein, such deviations from the theoretical predictions are attributed to corrections due to solvent and mode-independent internal friction.^62,63^ However, such a description for densely populated global minima (C type conformations) would imply an internal friction significantly larger than the solvent induced friction.^18^ Therefore, internal friction no longer remains a correction term, becoming the most dominant part. In addition, we show below that the different segments of the protein have distinct dynamical behavior, so that the averages on the scale of the whole chain are difficult to interpret.

We now proceed to examine the relaxation time spectra specific to the three conventional segments of the protein, see Fig. 6(b-d). For conformational class C, relaxation times for different modes show almost no separation for at *N*-(Fig. 6b) and *C*-terminal (Fig. 6d). For this type of conformations, the separation increases for the inner segment (Fig. 6c). Whereas, E and I type conformations display steeper decay profiles for all the segments apart from the hydrophobic *C*-terminal. Such spectra reflect a higher degree of internal fluctuations in these two configurational classes. However, the decay of *τ*_*p*_ with increasing *p* is still slower than the Zimm prediction.

Across all the types of conformations, hydrophobic *C*-terminal (Fig. 6d) exhibit relatively flatter spectra. The spectra indicates a collapsed structure with suppressed internal fluctuations. The observation is consistent with our previous estimates of *ν* for this segment, which showed globular signature across different classes of conformations. Thus, in addition to the heterogeneity for the same sequence at different regions of the conformational ensemble, we find chemically induced heterogeneity in internal fluctuations of different segments of the protein.

## Conclusions

Instead of a well-defined three dimensional structure, IDPs can access a wide range of conformations.^1^ The structural characterizations often employ static polymer scaling laws with effective scaling exponents.^7–10^ However, the presence of multiple quasi-stable conformations may challenge such descriptions.^13–15^ Different parts of conformational space can drastically differ in scaling behavior, effectively behaving like different polymers. In addition, different segments of the chain can, within a given class, have significantly different static and dynamic properties.

To investigate these issues, we sample the conformational space of a 209 residue long disordered *β*-casein using all-atom HREMD simulations^9,33,34^ in explicit solvent. AMBER99SB-ILDN force field along with explicit TIP4P-D water.^37,38^ We locate two shallow minima along with a global minimum. Associated structures can be classified into three classes, C, I and E with the single reaction coordinate *R*_*g*_. C type conformations are smallest in size and correspond to the global minimum. Whereas, those in class E are the largest and class I conformations have intermediate *R*_*g*_ values. E and I type structures are associated with the two local minima. Our global and local structural characterizations indicate that conformations belonging to C class are almost spherical, and compact. Conversely, E type conformations appear to behave like a polymer coil. I type conformations show intermediary features.

The existence of diverse quasi stable structures calls for separate investigations on scaling behaviors of each group of conformations. We indeed observe widely different Flory exponent (*ν*)^4,5^ for different conformational classes. As a result, there are differences in the scaling of Rouse mode amplitudes as well. ^64^ Both the static polymer scaling indicates that C type conformations are globular. Conversely, we find polymer coil-like features for conformational class E. Structures in class I have intermediate compactness. Therefore, in different regions of the conformational space, the IDP has structures akin to different kinds of polymers. We also investigate the differences in scaling features of the conventional segments of block-copolymer like *β*-casein.^24^ The hydrophobic *C*-terminal, across different classes, remain relatively collapsed. In contrast, the hydrophilic *N*-terminal exhibit a significant degree of swelling for all conformational types. Pronounced differences in compactness is observed for the inner segment. For class E, this segment is extremely expanded, but for class C it is much more compact. Once again, an intermediate level of swelling is noted for conformations in group I.

While conformational heterogeneity is known to be a characteristic property of an IDP, its implications on equilibrium dynamics remain poorly explored. We investigate the translational and rotational diffusion of the IDP using trajectories within funnels belonging to classes C, I, and E. Depending on the locality of the conformation, it can diffuse like a globule or a coil. We also show that the structural heterogeneity introduces heterogeneous internal fluctuation dynamics, which we elucidate probing the relaxation time of Rouse modes ^4,5,66^ corresponding to different segment lengths. Dynamics around global minimum yield an almost rigid-body like, flat spectrum, indicating insignificant internal fluctuations. Therefore, the dominant mode of relaxation is global rotation. We find comparatively steeper spectra for I and E type conformations. Yet, the decay profile is slower than the theoretical prediction. This can be attributed to solvent independent internal friction.^63^ The chemical segments of *β*-casein also show heterogeneous internal dynamics, with hydrophobic *C*-terminal having weak internal fluctuations. Whereas the two other domains show higher internal dynamics, especially in the extended conformation class. Therefore, our static and dynamic scaling analyses confirms the relevance of the conventional chemical segments of *β*-casein.

Our results emphasize the importance of population specific investigation for an appropriate characterization of the IDP. Our work also show that conformational variation can induce dynamic heterogeneity. However, different components of the motion, *e*.*g*., translational, rotational and internal can have different degree of heterogeneity. Therefore, all modes of the dynamics are to be considered to understand the relation between structure and dynamics in IDPs. Moreover, the investigation highlights polymeric principles to discern different segments of chemically heterogeneous block-copolymer like IDPs.

## Supporting Information Available

Convergence of HREMD simulations; comparison of experimental and simulation generated SAXS profiles; relaxation times for the Rouse modes corresponding to different conformational classes and their different segments.

## Acknowledgement

This work was supported by the ANR-DFG PRCI project ANR-21-CE06-0047. Numerical simulations were performed on GPU-accelerated partitions of the Jean Zay supercomputer hosted by GENCI-IDRIS (project AD010914227).

## References

(1) van der Lee, R. et al. Classification of Intrinsically Disordered Regions and Proteins. Chem. Rev. 2014, 114, 6589–6631.

(2) Wright, P. E.; Dyson, H. J. Intrinsically disordered proteins in cellular signalling and regulation. Nat. Rev. Mo. Cell Biol. 2015, 16, 18–29.

(3) Uversky, V. N.; Davé, V.; Iakoucheva, L. M.; Malaney, P.; Metallo, S. J.; Pathak, R. R.; Joerger, A. C. Pathological Unfoldomics of Uncontrolled Chaos: Intrinsically Disordered Proteins and Human Diseases. Chem. Rev. 2014, 114, 6844–6879.

(4) de Gennes, P. Scaling concepts in polymer physics; Cornell Univ. Pr., 1979; p 324 S.

(5) Doi, M.; Edwards, S. The Theory of Polymer Dynamics; Clarendon Press, 1986.

(6) Chan, H. S.; Dill, K. A. Polymer Principles in Protein Structure and Stability. Annual Review of Biophysics 1991, 20, 447–490.

(7) Petridis, L.; Schulz, R.; Smith, J. C. Simulation Analysis of the Temperature Dependence of Lignin Structure and Dynamics. J. Am. Chem. Soc. 2011, 133, 20277–20287.

(8) Hofmann, H.; Soranno, A.; Borgia, A.; Gast, K.; Nettels, D.; Schuler, B. Polymer scaling laws of unfolded and intrinsically disordered proteins quantified with single-molecule spectroscopy. Proc. Natl. Acad. Sci. 2012, 109, 16155–16160.

(9) Shrestha, U. R.; Juneja, P.; Zhang, Q.; Gurumoorthy, V.; Borreguero, J. M.; Urban, V.; Cheng, X.; Pingali, S. V.; Smith, J. C.; O’Neill, H. M.; Petridis, L. Generation of the configurational ensemble of an intrinsically disordered protein from unbiased molecular dynamics simulation. Proc. Natl. Acad. Sci. 2019, 116, 20446–20452.

(10) Huihui, J.; Ghosh, K. An analytical theory to describe sequence-specific inter-residue distance profiles for polyampholytes and intrinsically disordered proteins. The Journal of Chemical Physics 2020, 152, 161102.

(11) Brucale, M.; Schuler, B.; Samoři, B. Single-Molecule Studies of Intrinsically Disordered Proteins. Chem. Rev. 2014, 114, 3281–3317.

(12) Gomes, G.-N.; Gradinaru, C. C. Insights into the conformations and dynamics of intrinsically disordered proteins using single-molecule fluorescence. Biochim. Biophys. Acta - Proteins Proteom. 2017, 1865, 1696–1706.

(13) Firman, T.; Ghosh, K. Sequence charge decoration dictates coil-globule transition in intrinsically disordered proteins. J. Chem. Phys. 2017, 148, 123305.

(14) Sørensen, C. S.; Kjaergaard, M. Effective concentrations enforced by intrinsically disordered linkers are governed by polymer physics. Proc. Natl. Acad. Sci. 2019, 116, 23124–23131.

(15) Zeng, X.; Ruff, K. M.; Pappu, R. V. Competing interactions give rise to two-state behavior and switch-like transitions in charge-rich intrinsically disordered proteins. Proceedings of the National Academy of Sciences 2022, 119, e2200559119.

(16) Kundagrami, A.; Muthukumar, M. Effective Charge and CoilGlobule Transition of a Polyelectrolyte Chain. Macromolecules 2010, 43, 2574–2581.

(17) Qiao, Q.; Bowman, G. R.; Huang, X. Dynamics of an Intrinsically Disordered Protein Reveal Metastable Conformations That Potentially Seed Aggregation. J. Am. Chem. Soc. 2013, 135, 16092–16101.

(18) Ameseder, F.; Stingaciu, L. R.; Radulescu, A.; Holderer, O.; Falus, P.; Monkenbusch, M.; Biehl, R.; Richter, D.; Stadler, A. M. Structure and Dynamics of Intrinsically Disordered and Unfolded Proteins: Investigations using Small-Angle Scattering and Neutron Spin-Echo Spectroscopy. Biophys. J. 2019, 116, 490a–491a.

(19) Fischer, J.; Radulescu, A.; Falus, P.; Richter, D.; Biehl, R. Structure and Dynamics of Ribonuclease A during Thermal Unfolding: The Failure of the Zimm Model. J. Phys. Chem. B 2021, 125, 780–788.

(20) Henriques, J.; Arleth, L.; Lindorff-Larsen, K.; Skepö, M. On the Calculation of SAXS Profiles of Folded and Intrinsically Disordered Proteins from Computer Simulations. J. Mol. Biol. 2018, 430, 2521–2539, Intrinsically Disordered Proteins.

(21) Chan-Yao-Chong, M.; Durand, D.; Ha-Duong, T. Molecular Dynamics Simulations Combined with Nuclear Magnetic Resonance and/or Small-Angle X-ray Scattering Data for Characterizing Intrinsically Disordered Protein Conformational Ensembles. J. Chem. Inf. Model. 2019, 59, 1743–1758.

(22) Lincoff, J.; Sasmal, S.; Head-Gordon, T. The combined force field-sampling problem in simulations of disordered amyloid-β peptides. J. Chem. Phys. 2019, 150, 104108.

(23) Dumas, B. R.; Brignon, G.; Grosclaude, F.; Mercier, J.-C. Primary Structure of Bovine β casein: Complete sequence. Eur. J. Biochem. 1972, 25, 505–514.

(24) Farrell, H.; Wickham, E.; Unruh, J.; Qi, P.; Hoagland, P. Secondary structural studies of bovine caseins: temperature dependence of β-casein structure as analyzed by circular dichroism and FTIR spectroscopy and correlation with micellization. Food Hydrocoll. 2001, 15, 341–354.

(25) Zhou, M.; Xia, Y.; Cao, F.; Li, N.; Hemar, Y.; Tang, S.; Sun, Y. A theoretical and experimental investigation of the effect of sodium dodecyl sulfate on the structural and conformational properties of bovine β-casein. Soft Matter 2019, 15, 1551–1561.

(26) O’Connell, J.; Grinberg, V.; de Kruif, C. Association behavior of β-casein. Journal of Colloid and Interface Science 2003, 258, 33–39.

(27) Dauphas, S.; Mouhous-Riou, N.; Metro, B.; Mackie, A.; Wilde, P.; Anton, M.; Riaublanc, A. The supramolecular organisation of β-casein: effect on interfacial properties. Food Hydrocoll. 2005, 19, 387–393.

(28) Moitzi, C.; Portnaya, I.; Glatter, O.; Ramon, O.; Danino, D. Effect of Temperature on Self-Assembly of Bovine β-Casein above and below Isoelectric pH. Structural Analysis by Cryogenic-Transmission Electron Microscopy and Small-Angle X-ray Scattering. Langmuir 2008, 24, 3020–3029.

(29) Cragnell, C.; Choi, J.; Segad, M.; Lee, S.; Nilsson, L.; Skepö, M. Bovine β-casein has a polydisperse distribution of equilibrium micelles. Food Hydrocoll. 2017, 70, 65–68.

(30) Lin, Y.-H.; Chan, H. S. Phase Separation and Single-Chain Compactness of Charged Disordered Proteins Are Strongly Correlated. Biophys. J. 2017, 112, 2043–2046.

(31) Dignon, G. L.; Zheng, W.; Best, R. B.; Kim, Y. C.; Mittal, J. Relation between singlemolecule properties and phase behavior of intrinsically disordered proteins. Proc. Natl. Acad. Sci. 2018, 115, 9929–9934.

(32) Sugita, Y.; Okamoto, Y. Replica-exchange molecular dynamics method for protein folding. Chemical Physics Letters 1999, 314, 141–151.

(33) Wang, L.; Friesner, R. A.; Berne, B. J. Replica Exchange with Solute Scaling: A More Efficient Version of Replica Exchange with Solute Tempering (REST2). The Journal of Physical Chemistry B 2011, 115, 9431–9438.

(34) Bussi, G. Hamiltonian replica exchange in GROMACS: a flexible implementation. Molecular Physics 2014, 112, 379–384.

(35) Shrestha, U. R.; Smith, J. C.; Petridis, L. Full structural ensembles of intrinsically disordered proteins from unbiased molecular dynamics simulations. Commun. Biol. 2021, 4, 243.

(36) Yang, J.; Yan, R.; Roy, A.; Xu, D.; Poisson, J.; Zhang, Y. The I-TASSER Suite: protein structure and function prediction. Nat. Methods 2015, 12, 7–8.

(37) Lindorff-Larsen, K.; Piana, S.; Palmo, K.; Maragakis, P.; Klepeis, J. L.; Dror, R. O.; Shaw, D. E. Improved side-chain torsion potentials for the Amber ff99SB protein force field. Proteins 2010, 78, 1950–1958.

(38) Piana, S.; Donchev, A. G.; Robustelli, P.; Shaw, D. E. Water Dispersion Interactions Strongly Influence Simulated Structural Properties of Disordered Protein States. J. Phys. Chem. B 2015, 119, 5113–5123.

(39) Morozova, T. I.; García, N. A.; Barrat, J.-L. Temperature dependence of thermodynamic, dynamical, and dielectric properties of water models. J. Chem. Phys. 2022, 156, 126101.

(40) Henriques, J.; Skepö, M. Molecular Dynamics Simulations of Intrinsically Disordered Proteins: On the Accuracy of the TIP4P-D Water Model and the Representativeness of Protein Disorder Models. Journal of Chemical Theory and Computation 2016, 12, 3407–3415.

(41) Morozova, T. I.; García, N. A.; Matsarskaia, O.; Roosen-Runge, F.; Barrat, J.-L. Structural and Dynamical Properties of Elastin-Like Peptides near Their Lower Critical Solution Temperature. Biomacromolecules 2023, 24, 1912–1923.

(42) Shabane, P. S.; Izadi, S.; Onufriev, A. V. General Purpose Water Model Can Improve Atomistic Simulations of Intrinsically Disordered Proteins. J. Chem. Theory Comput. 2019, 15, 2620–2634.

(43) Bonomi, M.; Branduardi, D.; Bussi, G.; Camilloni, C.; Provasi, D.; Raiteri, P.; Donadio, D.; Marinelli, F.; Pietrucci, F.; Broglia, R. A.; Parrinello, M. PLUMED: A portable plugin for free-energy calculations with molecular dynamics. Comput. Phys. Commun. 2009, 180, 1961–1972.

(44) Bullerjahn, J. T.; von Bülow, S.; Heidari, M.; Hnin, J.; Hummer, G. Unwrapping NPT Simulations to Calculate Diffusion Coefficients. J. Chem. Theory Comput. 2023, 19, 3406–3417.

(45) Berendsen, H.; van der Spoel, D.; van Drunen, R. GROMACS: A message-passing parallel molecular dynamics implementation. Comp. Phys. Commun. 1995, 91, 43–56.

(46) Michaud-Agrawal, N.; Denning, E. J.; Woolf, T. B.; Beckstein, O. MDAnalysis: A toolkit for the analysis of molecular dynamics simulations. Journal of Computational Chemistry 2011, 32, 2319–2327.

(47) Efron, B. The Jackknife, the bootstrap and other resampling plans; Regional Conference Series in applied mathematics 38; Society for Industrial and applied mathematics: Philadelphia, Pa., 1982.

(48) Tavernelli, I.; Cotesta, S.; Di Iorio, E. E. Protein Dynamics, Thermal Stability, and Free-Energy Landscapes: A Molecular Dynamics Investigation. Biophys. J. 2003, 85, 2641–2649.

(49) Chebaro, Y.; Ballard, A. J.; Chakraborty, D.; Wales, D. J. Intrinsically Disordered Energy Landscapes. Sci. Rep. 2015, 5, 10386.

(50) Granata, D.; Baftizadeh, F.; Habchi, J.; Galvagnion, C.; De Simone, A.; Camilloni, C.; Laio, A.; Vendruscolo, M. The inverted free energy landscape of an intrinsically disordered peptide by simulations and experiments. Sci. Rep. 2015, 5, 15449.

(51) Ruskamo, S.; Chukhlieb, M.; Vahokoski, J.; Bhargav, S. P.; Liang, F.; Kursula, I.; Kursula, P. Juxtanodin is an intrinsically disordered F-actin-binding protein. Scientific Reports 2012, 2, 899.

(52) Rudnick, J.; Gaspari, G. The aspherity of random walks. Journal of Physics A: Mathematical and General 1986, 19, L191.

(53) Aronovitz, J.A.,; Nelson, D.R., Universal features of polymer shapes. J. Phys. France 1986, 47, 1445–1456.

(54) Dima, R. I.; Thirumalai, D. Asymmetry in the Shapes of Folded and Denatured States of Proteins. The Journal of Physical Chemistry B 2004, 108, 6564–6570.

(55) Rudnick, J.; Gaspari, G. The Shapes of Random Walks. Science 1987, 237, 384–389.

(56) Kirkwood, J. G.; Riseman, J. The Intrinsic Viscosities and Diffusion Constants of Flexible Macromolecules in Solution. The Journal of Chemical Physics 1948, 16, 565–573.

(57) Dünweg, B.; Kremer, K. Molecular dynamics simulation of a polymer chain in solution. The Journal of Chemical Physics 1993, 99, 6983–6997.

(58) Liu, B.; Dünweg, B. Translational diffusion of polymer chains with excluded volume and hydrodynamic interactions by Brownian dynamics simulation. The Journal of Chemical Physics 2003, 118, 8061–8072.

(59) Pesce, F.; Newcombe, E. A.; Seiffert, P.; Tranchant, E. E.; Olsen, J. G.; Grace, C. R.; Kragelund, B. B.; Lindorff-Larsen, K. Assessment of models for calculating the hydrodynamic radius of intrinsically disordered proteins. Biophys. J. 2023, 122, 310–321.

(60) Torricella, F.; Tugarinov, V.; Clore, G. M. Nucleation of Huntingtin Aggregation Proceeds via Conformational Conversion of Pre-Formed, Sparsely-Populated Tetramers. Adv. Sci. 2024, 11, 2309217.

(61) Pounot, K.; Piersson, C.; Goring, A. K.; Rosu, F.; Gabelica, V.; Weik, M.; Han, S.; Fichou, Y. Mutations in Tau Protein Promote Aggregation by Favoring Extended Conformations. JACS Au 2024, 4, 92–100, extended conformations have shorter lag time, in other words more prone to aggregation.

(62) Soranno, A.; Buchli, B.; Nettels, D.; Cheng, R. R.; Müller-Späth, S.; Pfeil, S. H.; Hoffmann, A.; Lipman, E. A.; Makarov, D. E.; Schuler, B. Quantifying internal friction in unfolded and intrinsically disordered proteins with single-molecule spectroscopy. Proc. Natl. Acad. Sci. 2012, 109, 17800–17806.

(63) Khatri, B. S.; McLeish, T. C. B. Rouse Model with Internal Friction: A Coarse Grained Framework for Single Biopolymer Dynamics. Macromolecules 2007, 40, 6770–6777.

(64) Földes, T.; Lesage, A.; Barbi, M. Assessing the Polymer Coil-Globule State from the Very First Spectral Modes. Phys. Rev. Lett. 2021, 127, 277801.

(65) Panja, D.; Barkema, G. T. Rouse modes of self-avoiding flexible polymers. J. Chem. Phys. 2009, 131, 154903.

(66) Khokhlov, A. Statistical Physics of Macromolecules; AIP series in polymers and complex materials; AIP Press, 1994.

(67) Yeh, I.-C.; Hummer, G. System-Size Dependence of Diffusion Coefficients and Viscosities from Molecular Dynamics Simulations with Periodic Boundary Conditions. J. Phys. Chem. B 2004, 108, 15873–15879.

(68) Linke, M.; Köfinger, J.; Hummer, G. Rotational Diffusion Depends on Box Size in Molecular Dynamics Simulations. J. Phys. Chem. Lett. 2018, 9, 2874–2878.

(69) Zimm, B. H. Dynamics of Polymer Molecules in Dilute Solution: Viscoelasticity, Flow Birefringence and Dielectric Loss. J. Chem. Phys. 1956, 24, 269–278.

