## Supplemental figures 1-4 for "Intrinsically disordered proteins can behave as different polymers across their conformational ensemble"

### Intrinsically Disordered Protein can Behave as Different Polymers in Different Regions of the Conformational Ensemble

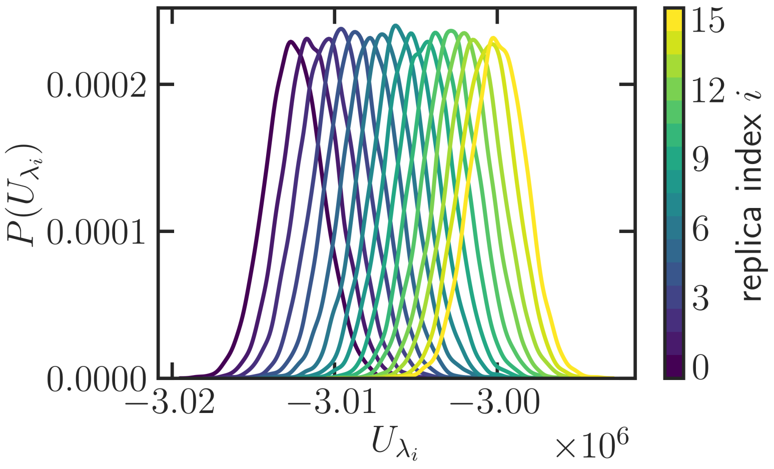


Figure 1: Overlap of potential energy distribution for neighboring replicas in Hamiltonian replica exchange simulations indicate sufficiency of the number of replicas and absence of any phase transitions. Average exchange probability between the replicas is $\approx 0.21$.

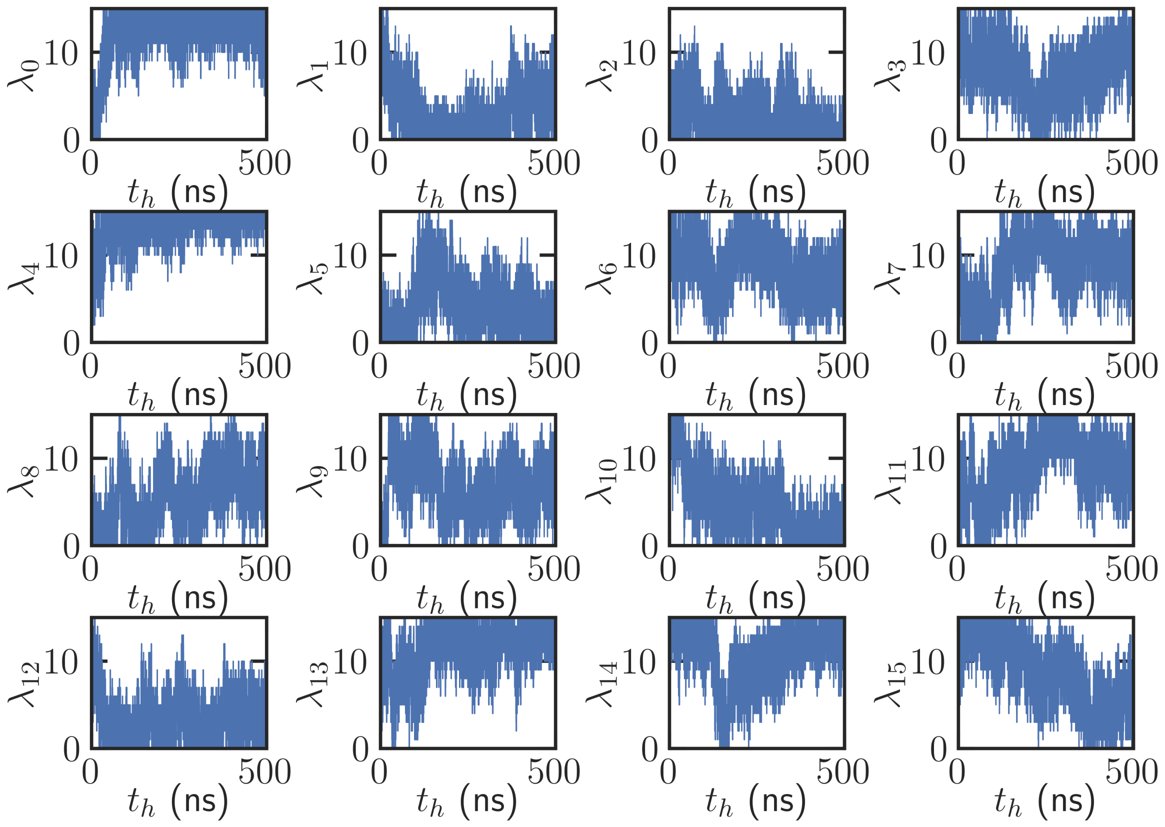


Figure 2: Trajectories of the conformations in the replica index space in the HREMD simulations. Each of the 16 replica passes through all the indices between $[0, 15]$ reflecting convergence of the simulations.

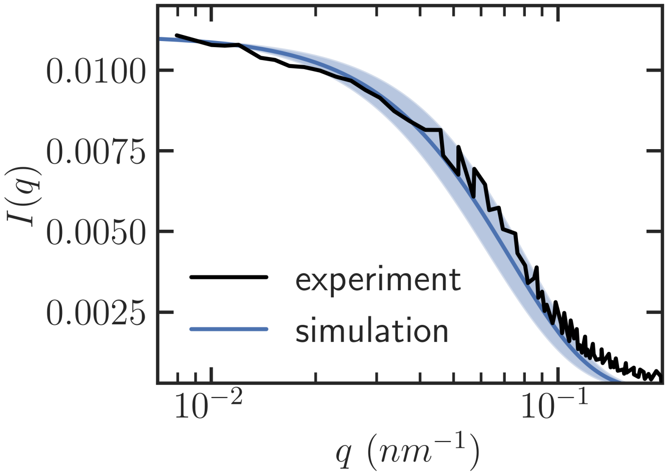


Figure 3: Experimental small angle X-ray scattering (SAXS) data obtained digitizing Fig. 2A in Ref. 1. Zhou *et. al.* performed the measurements at pH 9.0. Alongside, SAXS profile generated with conformations in Hamiltonian replica exchange molecular dynamics simulations (HREMD) simulations using CRYSOL. Consistency between the experiments and simulations in small $q$ regime confirms appropriate sampling of the radius of gyration, $R_g$. Mismatch between the two plots appear at length scales comparable to few $C_\alpha$ atoms. This may be because of lower fraction of order in simulated conformations. The higher pH value in experiments can also be the reason behind the inconsistency.

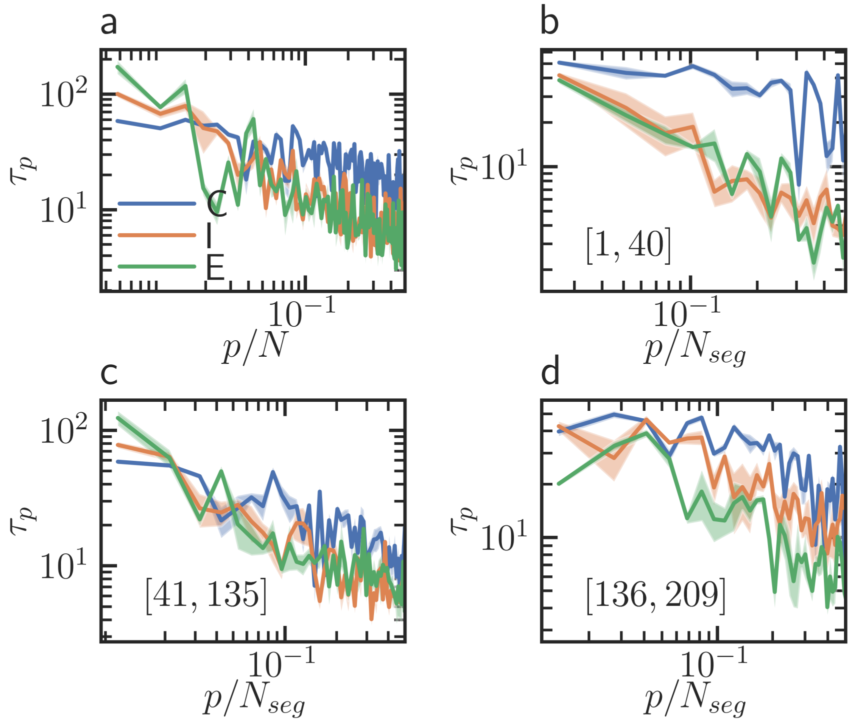


Figure 4: Relaxation time $\tau_p$ associated with the $p$th Rouse modes. The frames are same as Figure 6 in main text. However, $\tau_p$s are not scaled by $\tau_1$. Results are presented for the whole chain (a) and for the conventional segments (b-d) discussed in the main text.

### References

(1) Zhou, M.; Xia, Y.; Cao, F.; Li, N.; Hemar, Y.; Tang, S.; Sun, Y. A theoretical and experimental investigation of the effect of sodium dodecyl sulfate on the structural and conformational properties of bovine $\beta$-casein. *Soft Matter* **2019**, *15*, 1551–1561.
